# Detection of extended-spectrum beta-lactamase (ESBL) genes and plasmid replicons in Enterobacteriaceae using PlasmidSPAdes assembly of short-read sequence data

**DOI:** 10.1101/863316

**Authors:** Joep J.J.M. Stohr, Marjolein F.Q. Kluytmans-van den Bergh, Ronald Wedema, Alexander W. Friedrich, Jan A.J.W. Kluytmans, John W.A. Rossen

**Affiliations:** Department of Infection Control, Amphia Hospital, Breda, the Netherlands; Laboratory for Medical Microbiology and Immunology, Elisabeth-TweeSteden Hospital, Tilburg, the Netherlands; Amphia Academy Infectious Disease Foundation, Amphia Hospital, Breda, the Netherlands; Julius Center for Health Sciences and Primary Care, University Medical Center Utrecht, Utrecht, the Netherlands; Department of Life Science and Technology, Hanze University of Applied Sciences, Groningen, the Netherlands; Department of Medical Microbiology and Infection Prevention, University of Groningen, University Medical Center Groningen, Groningen, the Netherlands

## Abstract

**INTRODUCTION:** Knowledge of the epidemiology of plasmids is essential for understanding the evolution and spread of antimicrobial resistance. PlasmidSPAdes attempts to reconstruct plasmids using short-read sequence data. Accurate detection of extended-spectrum beta-lactamase (ESBL) genes and plasmid replicon genes is a prerequisite for the use of plasmid assembly tools to investigate the role of plasmids in the spread and evolution of ESBL production in Enterobacteriaceae. This study evaluated the performance of PlasmidSPAdes plasmid assembly for Enterobacteriaceae in terms of detection of ESBL-encoding genes, plasmid replicons and chromosomal genes, and assessed the effect of k-mer size.

**METHODS:** Short-read sequence data of 59 ESBL-producing Enterobacteriaceae were assembled with PlasmidSPAdes using different k-mer sizes (21, 33, 55, 77, 99 and 127). For every k-mer size, the presence of ESBL genes, plasmid replicons and a selection of chromosomal genes in the plasmid assembly was determined.

**RESULTS:** Out of 241 plasmid replicons and 66 ESBL genes detected by whole-genome assemblies, 213 plasmid replicons (88%; 95% Confidence Interval (CI): 83.9-91.9) and 43 ESBL genes (65%; 95%CI: 53.1-75.6) were detected in the plasmid assemblies obtained by PlasmidSPAdes. For most ESBL genes (83.3%) and plasmid replicons (72.0%), detection results did not differ between the k-mer sizes used in the plasmid assembly. No optimal k-mer size could be defined for the number of ESBL genes and plasmid replicons detected. For most isolates, the number of chromosomal genes detected in the plasmid assemblies decreased with increasing k-mer size.

**CONCLUSION:** Based on our findings, PlasmidSPAdes is not a suitable plasmid assembly tool for short-read sequence data of ESBL-encoding plasmids of Enterobacteriaceae.

## Introduction

Plasmids are small DNA molecules that naturally exist within bacterial cells and replicate independently from the chromosome (^1^). They can harbour genes involved in virulence and antibiotic resistance and are important vectors for horizontal gene transfer (^2,3^). Knowledge of the epidemiology of plasmids is essential for understanding the evolution and spread of antimicrobial resistance. It remains challenging to study plasmids using short-read whole-genome sequencing (WGS) data (^4^). Repeated sequences, often shared between plasmid and chromosomal DNA, hinder the assembly of the bacterial genome from short-read sequence data, often resulting in contigs of which the origin, either plasmid or chromosomal, cannot be resolved. In recent years, several tools to study plasmids using WGS data were developed and include PLACNET, PlasmidFinder, cBar, HyAsP, MOB-suite, PlasmidTron, Recycler, and PlasmidSPAdes (^4–11^). Only Recycler and PlasmidSPAdes are fully automated computer programs that aim to *de novo* reconstruct whole plasmid sequences from short-read sequence data (^7,8^). Recent studies benchmarked the PlasmidSPAdes and Recycler algorithm for genomes of Gram-negative bacteria (^12,13^). These studies showed PlasmidSPAdes outperforming Recycler in terms of the plasmid and chromosomal fraction detected in the putative plasmids created by these algorithms (^12^). However, data on the performance of PlasmidSPAdes in representative and well-defined clinical data sets of Enterobacteriaceae remain limited (^11,13^). PlasmidSPAdes uses the SPAdes *De Bruijn* graph assembler (^8^), in which k-mer size is an important parameter (^14–16^) that influences contig size, entanglement and location of contigs in the assembly graph, and accuracy and coverage of the assembly (^13^). An algorithm optimizing the k-mer size selection for whole-genome assemblies was previously published (^16^). Yet, no data are available on how this key setting affects the performance of plasmid assemblies using PlasmidSPAdes. The objective of this study was to evaluate the performance of PlasmidSPAdes plasmid assembly of short-read sequence data of extended-spectrum beta-lactamase-producing Enterobacteriaceae (ESBL-E) in terms of detection of ESBL-encoding genes, plasmid replicons and chromosomal genes, and to assess the effect of k-mer size.

## Methods

### Selection of whole-genome sequencing data

Raw whole-genome sequencing reads of 59 ESBL-E isolates were selected from study accession number PREJB15226 of the publicly available European Nucleotide Archive (ENA) of the European Bioinformatics Institute (EBI) (^17^) and included 21 *Escherichia coli*, 10 *Klebsiella pneumoniae*, 10 *Klebsiella oxytoca*, 10 *Enterobacter cloacae complex* and 8 *Citrobacter* spp. The selection included a variety of plasmid types. Sequence data in the database were generated on either an Illumina MiSeq sequencer (Illumina, San Diego, CA, USA) or a HiSeq 2500 sequencer (Illumina, San Diego, CA, USA).

### *De novo* whole-genome assembly (WGA)

*De novo* WGA was performed using CLC Genomics Workbench 7.0.4 (Qiagen, Hilden, Germany) (^17^). The k-mer size used in the assembly was based on the maximum N50 value (the largest scaffold length, N, such that 50% of the genome size is made up of scaffolds with a length of at least N) (^17^).

### *De novo* plasmid assembly

*De novo* plasmid assembly with raw read error correction was performed using PlasmidSPAdes version 3.9 with default settings. PlasmidSPAdes uses the SPAdes whole-genome assembler to construct an assembly graph and subsequently excludes chromosomal edges from the assembly graph based on k-mer coverage and edge size. The algorithm calculates the median k-mer coverage of all long edges (>10 kilobases (kb)). Then, long edges are classified as chromosomal when the k-mer coverage of the edge divided by the median k-mer coverage of all long edges is higher than 1 minus the *maxDeviation* parameter (default: 0.3), but lower than 1 plus the *maxDeviation* parameter. Long chromosomal edges are iteratively removed from the assembly graph unless they belong to a connected component with no dead-end edges or to a component with a size below the *maxComponentSize* (default: 150kb). When at least one long edge is removed during the previous step, all short dead-end edges are removed. An edge (*a,b*) is considered a dead-end edge when either node *a* has indegree zero or node *b* has outdegree zero but not both. After these iterative exclusion steps, all non-plasmidic connected components are excluded from the assembly graph. A component is classified as plasmidic when it is composed of a single-loop edge of at least *minCirc* (default: 1kb) or if it exceeds *minCompsize* (default: 10kb) (^8^). Incremental plasmid assemblies were performed at different k-mer sizes. Both MiSeq and HiSeq 2500 derived reads were assembled at k-mer sizes 21, 33, 55 and 77. The increased read length for MiSeq derived reads enabled additional assembly of MiSeq derived reads at k-mer sizes 99 and 127. For each k-mer size, the plasmid assembly and the assembled genome before chromosomal removal were retained. Quality control statistics were generated for all plasmid assemblies using Quast v.5.0.2. (^18^). The median k-mer coverages of all long edges (>10kb) in the assembly as defined by Antipov *et al*. (^8^) was calculated for each PlasmidSPAdes assembly using a custom script.

### Detection of ESBL genes and plasmid replicons

The CLC derived WGAs and the PlasmidSPAdes plasmid assemblies and assembled genomes before chromosomal removal were uploaded onto the online bioinformatics tools ResFinder v2.1 and PlasmidFinder v.1.3.1 (Center for Genetic Epidemiology, DTU, Denmark) (^5,19^). ESBL genes were called when at least 60% of the sequence length of the gene in the ResFinder database was covered with a sequence identity of at least 90%. Plasmid replicon genes were called when at least 60% of the sequence length of the replicon gene in the PlasmidFinder database was covered with a sequence identity of at least 80%. ESBL genes and plasmid replicons that were called for the plasmid assemblies were compared with those called for the corresponding WGAs. If genes were detected more than once within the plasmid assembly, only one of the called genes was used for the comparison. ESBL genes and plasmid replicons that were called for the plasmid assembly but not for the WGA were classified as additionally found ESBL genes or plasmid replicons unless the ESBL gene or plasmid replicon was from the same resistance class (e.g. *bla*_CTX-M_, *bla*_SHV_ or *bla*_TEM_) or plasmid incompatibility group but with a lower ambiguity score. The ambiguity score is defined as the percentage of the sequence length that is aligned with the called gene multiplied by the sequence identity of this alignment (^20^).

### Detection of chromosomal DNA

Chromosomal genes were blasted against the plasmid assemblies using BLAST+ from ABRicate version 0.8.2 (https://github.com/tseemann/abricate) to assess non-specific incorporation of chromosomal DNA in the plasmid assemblies. Chromosomal genes were derived from recently developed species-specific whole-genome multilocus sequence typing (wgMLST) schemes that excluded plasmid sequences (^17^). Similar to the settings used for creating the wgMLST schemes, wgMLST genes were called when 100% of the sequence length was covered with an identity of at least 90% (^17^).

### Assessment of coverage deviation

For all ESBL gene-or plasmid replicon-containing contigs in the PlasmidSPAdes derived assembly before chromosome removal, the k-mer coverage was compared with the median k-mer coverage of all long edges (>10kb) in the assembly. Only contigs containing an ESBL gene and/or plasmid replicon that was also detected in the corresponding CLC-derived WGA were included in the analysis. A contig k-mer coverage below 0.7 or above 1.3 times the median k-mer coverage of all long edges (>10kb) in the assembly was defined as deviating. Subsequently, all contigs containing an ESBL-gene or plasmid replicon with and without a deviating coverage were checked for their presence in the plasmid assemblies. For each assembly that included a contig with a non-deviating k-mer coverage, the assembly graph before chromosomal removal was inspected using BANDAGE (^21^). As a sensitivity analysis, the total number of long contigs and the ESBL gene-and plasmid replicon-containing contigs with a k-mer coverage below 0.8 or above 1.2 the median k-mer coverage was calculated to assess the effect of reducing the *maxDeviation* parameter from 0.3 to 0.2.

## Results

PlasmidSPAdes assembly characteristics before and after chromosome removal were dependent on k-mer size (**Supplementary table 1**). An increasing k-mer size was associated with a decrease in the median k-mer coverage and the number of contigs but with an increase in the N50 and the maximum contig size. The size of the plasmid assemblies was largest for the smallest k-mer size, decreased with increasing k-mer size, but did not further decrease at k-mer sizes higher than 55. For every k-mer size used the median size of the plasmid assembly was larger than the *maxComponent* size parameter of 150kb.

For 59 isolates, 241 plasmid replicons and 66 ESBL genes were detected in the WGA. Of those, 213 plasmid replicons (88%; 95% confidence interval (CI): 83.7-91.9) and 43 ESBL genes (65%; 95% CI: 53.1-75.6) were detected in at least one of the plasmid assemblies (**Table 1**). Eight plasmid replicons and one ESBL gene that were detected in the plasmid assembly were not identified in the WGA, despite their presence in the raw reads (**Supplementary table 2**).

**Table 1.** Detection of ESBL genes and plasmid replicons in the plasmid assembly as compared to the whole genome assembly.

|  | Gene/replicon detected in whole-genome assembly | Gene/replicon detected in at least one of the plasmid assemblies |  |
| --- | --- | --- | --- |
|  | n | n | % |
| <u>ESBL gene</u> |  |  |  |
| *bla*<sub>CTX-M-1</sub> | 4 | 4 | 100 |
| *bla*<sub>CTX-M-3</sub> | 1 | 1 | 100 |
| *bla*<sub>CTX-M-9</sub> | 16 | 4 | 25 |
| *bla*<sub>CTX-M-14</sub> | 4 | 4 | 100 |
| *bla*<sub>CTX-M-14b</sub> | 1 | 1 | 100 |
| *bla*<sub>CTX-M-15</sub> | 17 | 14 | 82 |
| *bla*<sub>CTX-M-27</sub> | 1 | 1 | 100 |
| *bla*<sub>CTX-M-30</sub> | 2 | 2 | 100 |
| *bla*<sub>CTX-M-55</sub> | 1 | 1 | 100 |
| *bla*<sub>GES-1</sub> | 1 | 0 | 0 |
| *bla*<sub>SHV-12</sub> | 13 | 8 | 62 |
| *bla*<sub>SHV-28</sub> | 1 | 0 | 0 |
| *bla*<sub>TEM-15</sub> | 1 | 0 | 0 |
| *bla*<sub>TEM-52B</sub> | 3 | 3 | 100 |
| Total | 66 | 43 | 65 |
| <u>Plasmid replicon family</u> |  |  |  |
| Col | 61 | 56 | 92 |
| IncA/C2 | 2 | 2 | 100 |
| IncB/O/K/Z | 4 | 4 | 100 |
| IncFIA | 12 | 9 | 75 |
| IncFIB | 42 | 38 | 90 |
| IncFII | 45 | 37 | 82 |
| IncHI2 | 34 | 27 | 79 |
| IncI1 | 10 | 10 | 100 |
| IncI2 | 1 | 1 | 100 |
| IncL/M | 1 | 1 | 100 |
| IncN | 2 | 2 | 100 |
| IncN3 | 2 | 2 | 100 |
| IncP1 | 1 | 0 | 0 |
| IncQ1 | 2 | 2 | 100 |
| IncQ2 | 1 | 1 | 100 |
| IncR | 5 | 5 | 100 |
| IncX1 | 8 | 7 | 88 |
| IncX3 | 4 | 3 | 75 |
| IncX4 | 3 | 3 | 100 |
| IncY | 4 | 3 | 75 |
| Total | 241 | 213 | 88 |

The detection of ESBL-genes and plasmid replicons was not dependent on k-mer size. Concordant results for all k-mer size plasmid assemblies were found for 173 (72%) of 241 plasmid replicons and 55 (83%) of 66 ESBL genes. For individual k-mer sizes, the percentage of plasmid replicons detected ranged from 72% (k-mer size 55) up to 81% (k-mer size 127) and the percentage of ESBL genes detected from 53% (k-mer size 21) up to 61% (k-mer sizes 55 and 77), with no statistically significant differences (**Supplementary table 3**). Thirteen plasmid replicons and two ESBL genes were detected at one k-mer size only, although at varying k-mer sizes.

The retrieval of plasmid replicons and ESBL genes differed between bacterial species. For plasmid replicons, retrieval ranged from 67% (14/21) in *Citrobacter* spp. to 96% (44/46) in *K. oxytoca*, and for ESBL genes from 17% (2/12) in *E. cloacae* to 91% (10/11) in *K. pneumoniae*) (**Supplementary table 3**). The retrieval of ESBL genes differed between plasmid families. For isolates with an IncHI2 replicon, e.g., only 5 of 22 (23%) WGA detected ESBL genes were detected in at least one of the plasmid assemblies as compared to 32 of 42 (76%) WGA detected ESBL genes in isolates with an IncFIB replicon. Only the IncQ2 plasmid replicon that was detected in the WGA of 1 isolate had a lower retrieval rate of ESBL genes in its corresponding isolate (**Supplementary table 4**).

In general, the percentage of wgMLST genes detected in the plasmid assemblies decreased with increasing k-mer size (**Figure 1**). In several *K. oxytoca* isolates and one *E. coli* isolate, however, the percentage of wgMLST genes detected increased when assembling with a larger k-mer size.

**Figure 1.**
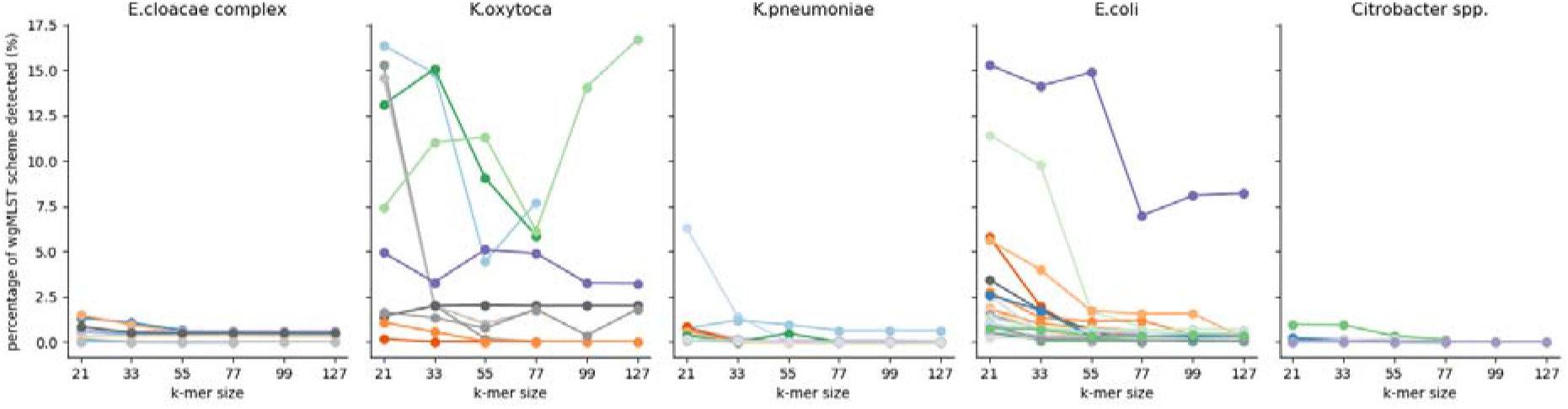
The percentage of wgMLST genes that were detected in the plasmid assemblies. Every line represents a plasmid assembly of an isolate.

A total number of 312 ESBL-gene-containing contigs and 989 plasmid replicon-containing contigs were present in any of the assembly graphs before removal of the bacterial chromosome. Of those, 128 (41%) of ESBL-gene containing contigs and 192 (19%) plasmid containing contigs were not present in the plasmid assemblies. The main reason for being excluded from the assembly graph was a non-deviating k-mer coverage in 119 (38%) of ESBL-E containing contigs and 113 (11%) of plasmid replicon-containing contigs. A contig size smaller than 10 kb explained the exclusion of 8 (3%) ESBL-containing contigs and 76 (8%) plasmid containing contigs with deviating k-mer coverage, which probably has resulted in the removal of a ‘dead-end edge’ or ‘non-plasmidic component’. Four contigs containing an ESBL gene or plasmid replicon were not included in the plasmid assembly despite being longer than 10kb and having a coverage that deviated from the median k-mer coverage of the assembly.

Five ESBL genes-and sixteen plasmid replicons contigs that were included in the plasmid assembly but not located on a contig with a coverage that deviated from the median were part of a component with no dead-end edges smaller than *maxComponentSize* (<150kb) (**Table 2 and Supplementary figure 2**).

**Table 2.** The effect of deviating k-mer coverage on the presence of WGA detected ESBL gene containing contigs and plasmid replicon containing contigs in the plasmid assembly.

| Deviating coverage* | ESBL gene detected in WGA |  | Plasmid replicon detected in WGA |  |
| --- | --- | --- | --- | --- |
|  | Detected in | Not detected in | Detected in | Not detected in |
|  | plasmid | plasmid | plasmid | plasmid |
|  | assembly | assembly | assembly | assembly |
|  | n (%) | n (%) | n (%) | n (%) |
| Yes | 179 (95) | 9 (5) | 780 (91) | 79 (9) |
| No | 5 (4) | 119 (96) | 17 (13) | 113 (87) |

\* Deviating coverage = k-mer coverage of contig/median coverage of all long edges (>10kb) > 1.3 or k-mer coverage of contig/median coverage of all long edges (>10kb) < 0.7

The percentage of contigs, encoding an ESBL gene or plasmid replicon, that were excluded from the assembly despite having a deviating coverage differed per k-mer size used in the plasmid assembly (**Supplementary table 5**).

The percentage of ESBL gene-and plasmid replicon-containing contigs with a deviating k-mer coverage increased when the *maxDeviation* parameter was reduced from 0.3 to 0.2 (**Table 3**). At the same time, the total number of long contigs (>10kb) with a deviating k-mer coverage also increased, which may reduce the specificity of the plasmidome.

**Table 3.**
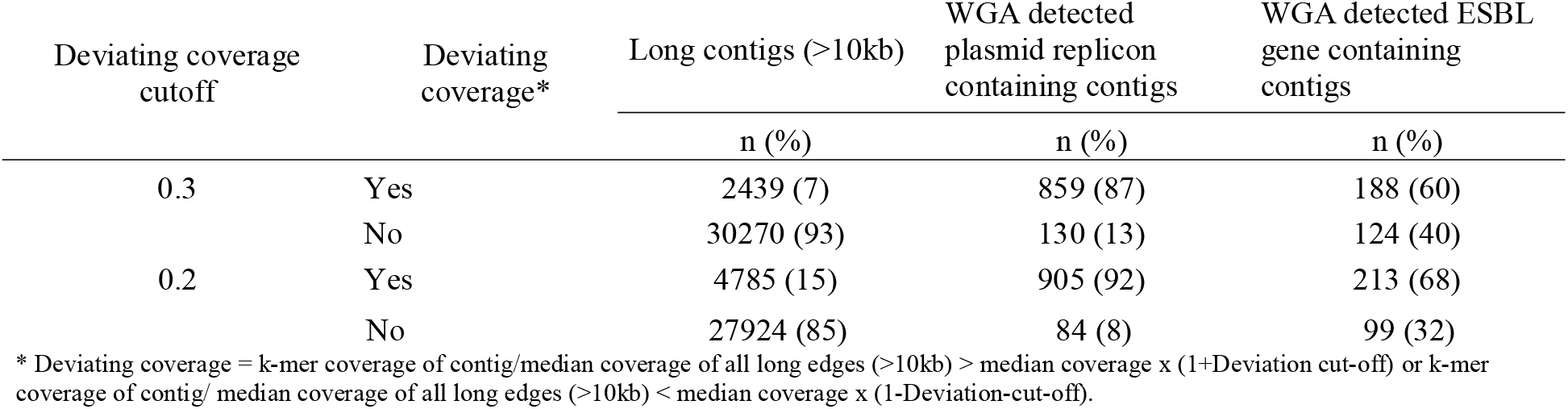
The effect of deviating k-mer coverage cut-off alteration on the number of long contigs (>10kb), ESBL gene-and plasmid replicon-containing contigs with a k-mer coverage that deviated from the median k-mer coverage of all long contigs (>10kb).

## Discussion

PlasmidSPAdes assembly was not useful to reliably detect the presence of ESBL-genes and plasmid replicons in short-read sequence data of ESBL-E, independent of k-mer size. A non-deviating k-mer coverage was the main reason for the exclusion of ESBL-gene-or plasmid replicon containing contigs from the plasmid assembly. The presence of chromosomal genes in the plasmid assembly varied between bacterial species and was not consistently related to k-mer size.

Retrieval rates of ESBL genes and plasmid replicons are similar to the retrieval rate of plasmid DNA by plasmidSPAdes in earlier studies (^10–12^), in which long-read sequence data were used as a reference for plasmid DNA detection. However, these studies included Enterobacteriaceae sequences belonging to different undefined datasets, used only the default setting of the plasmidSPAdes algorithm, and did not evaluate which steps in the plasmidSPAdes algorithm were critical in the plasmid assembly.

Although for most isolates the presence of chromosomal genes decreased with increasing k-mer size, in four *K. oxytoca* isolates and one *E. coli* isolate, several chromosomal contigs were not excluded from the plasmid assemblies at the large k-mer sizes. Apparently, large chromosomal contigs with a non-deviating k-mer coverage were formed at these large k-mer sizes, resulting in the erroneous assignment of these contigs as plasmidic.

The low detection rate of both plasmid replicons and ESBL genes in *Enterobacter* and *Citrobacter* isolates when compared to the other genera investigated coincides with a low detection rate of the IncHI2a and *bla*_CTX-M-9_ gene when compared to the other plasmid replicons and ESBL genes. This possibly is because, contrary to the isolates of the other investigated genera, half or more than half of the *Citrobacter* and *Enterobacter* isolates contained a *bla*_CTX-M-9_ and IncHI2a replicon in our selection. The co-existence of these IncHI2 plasmid replicons and *bla*_CTX-M-9_ genes in *E. cloacae complex* and *Citrobacter spp* was also seen in other studies (^2,22–24^).

The poor retrieval rates of ESBL genes and plasmid replicons in plasmidSPAdes derived plasmid assemblies confirms that ESBL carrying plasmids in Enterobacteriaceae frequently belong to plasmid families with copy numbers that resemble the chromosome (^1,2,25^). The higher retrieval rate for plasmid replicons than for ESBL genes may be explained by the presence of other (no ESBL gene-containing) plasmids with a higher copy number or a higher copy number of the plasmid replicon itself, increasing the k-mer coverage of the contig on which the plasmid replicon is located. Our findings suggest that lowering of the maxDeviation parameter may increase the retrieval rate of plasmid replicon-and ESBL gene-containing contigs, but may, on the other hand, reduce the specificity of the plasmid assembly. Since the *maxDeviation* parameter could not be adjusted in the command line of the PlasmidSPAdes version used in this study, the actual effect of the alteration of this parameter on the plasmid assemblies in our dataset remains unknown. The study of *Page et al*. also reported that read coverage can be a major determinant on PlasmidSPAdes performance (^9^). However, this study only included one genome, and read coverage alterations were manipulated *in silico* (^9^). Despite coverage information being the primary determinant for being included in the plasmid assemblies, several genes encoded on a contig with a deviating k-mer coverage were not incorporated. Most of these genes were located on contigs smaller than 10kb suggesting that either the “short dead-end edge” removal step or the “non-plasmidic component” removal step might falsely exclude these contigs with a deviating k-mer coverage from the plasmid assembly. Only a limited number of ESBL genes and plasmid replicons were incorporated in the plasmid assemblies when not present on a contig with a deviating k-mer coverage as defined by PlasmidSPAdes. Given that the median plasmid assembly size in our isolates was larger than 150kb for all the k-mer sizes used and given the possible entanglement of the various plasmids in one component, as observed in previous studies (^12^), increasing the *maxComponentSize* could include more genes through this “escape route” without incorporating more chromosomal DNA in the plasmid assemblies.

A strength of our study is that we studied a broad spectrum of ESBL-producing Enterobacteriaceae isolates, including 5 different genera of various sequence types, harbouring 21 different plasmid families. All isolates belong to a well-defined collection of ESBL-producing isolates that were collected, cultured and whole-genome sequenced using the same methods.

On the other hand, the use of plasmid replicon, ESBL gene and wgMLST gene detection instead of a complete genome as a reference to evaluate the plasmid assembly algorithm may have limited the resolution at which the algorithm could be evaluated. However, accurately detecting ESBL genes and plasmid replicon genes is a prerequisite for the use of plasmid assembly tools to investigate the role of plasmids in the spread and evolution of ESBL production in Enterobacteriaceae.

In conclusion, based on our data plasmidSPAdes is not a suitable plasmid assembly tool for short-read sequence data of ESBL-encoding plasmids of Enterobacteriaceae.

## Acknowledgments

We are grateful to the members of the SoM Study Group for their contribution to the collection, culturing and whole-genome sequencing of ESBL-E isolates.

## Supplementary materials

Supplementary tables 1,2,3,4,5.

Supplementary figures 1,2.

## References

1. Partridge SR, Kwong SM, Firth N, Jensen SO. Mobile genetic elements associated with antimicrobial resistance. Clin Microbiol Rev. 2018;31(4):1–61. doi:10.1128/CMR.00088-17

2. Carattoli A. Resistance plasmid families in Enterobacteriaceae. Antimicrob Agents Chemother. 2009;53(6):2227–2238. doi:10.1128/AAC.01707-08

3. Carattoli A. Plasmids and the spread of resistance. Int J Med Microbiol. 2013;303(6-7):298–304. doi:10.1016/j.ijmm.2013.02.001

4. Lanza VF, de Toro M, Garcillán-Barcia MP, et al. Plasmid Flux in Escherichia coli ST131 Sublineages, Analyzed by Plasmid Constellation Network (PLACNET), a New Method for Plasmid Reconstruction from Whole Genome Sequences. PLoS Genet. 2014;10(12). doi:10.1371/journal.pgen.1004766

5. Carattoli A, Zankari E, Garciá-Fernández A, et al. In Silico detection and typing of plasmids using plasmidfinder and plasmid multilocus sequence typing. Antimicrob Agents Chemother. 2014;58(7):3895–3903. doi:10.1128/AAC.02412-14

6. Zhou F, Xu Y. cBar: A computer program to distinguish plasmid-derived from chromosome-derived sequence fragments in metagenomics data. Bioinformatics. 2010;26(16):2051–2052. doi:10.1093/bioinformatics/btq299

7. Rozov R, Kav AB, Bogumil D, et al. Recycler: An algorithm for detecting plasmids from de novo assembly graphs. Bioinformatics. 2017;33(4):475–482. doi:10.1093/bioinformatics/btw651

8. Antipov D, Hartwick N, Shen M, Raiko M, Lapidus A, Pevzner PA. PlasmidSPAdes: Assembling plasmids from whole genome sequencing data. Bioinformatics. 2016;32(22):3380–3387. doi:10.1093/bioinformatics/btw493

9. Page AJ, Wailan A, Shao Y, et al. PlasmidTron: assembling the cause of phenotypes and genotypes from NGS data. Microb genomics. 2018;4(3):1–6. doi:10.1099/mgen.0.000164

10. Robertson J, Nash JHE. MOB-suite: software tools for clustering, reconstruction and typing of plasmids from draft assemblies. Microb genomics. 2018;4(8). doi:10.1099/mgen.0.000206

11. Müller R, Chauve C. HyAsP, a greedy tool for plasmids identification. Valencia A, ed. Bioinformatics. 2019;35(21):4436–4439. doi:10.1093/bioinformatics/btz413

12. Arredondo-Alonso S, Willems RJ, van Schaik W, Schürch AC. On the (im)possibility of reconstructing plasmids from whole-genome short-read sequencing data. Microb Genomics. 2017;(im). doi:10.1099/mgen.0.000128

13. Laczny CC, Galata V, Plum A, Posch AE, Keller A. Assessing the heterogeneity of in silico plasmid predictions based on whole-genome-sequenced clinical isolates. Brief Bioinform. 2019;20(3):857–865. doi:10.1093/bib/bbx162

14. Zerbino DR, Birney E. Velvet: Algorithms for de novo short read assembly using de Bruijn graphs. Genome Res. 2008;18(5):821–829. doi:10.1101/gr.074492.107

15. Nurk S, Bankevich A, Antipov D, et al. Assembling Genomes and Mini-metagenomes from Highly Chimeric Reads. In:; 2013:158–170. doi:10.1007/978-3-642-37195-0_13

16. Chikhi R, Medvedev P. Informed and automated k-mer size selection for genome assembly. Bioinformatics. 2014;30(1):31–37. doi:10.1093/bioinformatics/btt310

17. Kluytmans-van den Bergh MFQ, Rossen JWA, Bruijning-Verhagen PCJ, et al. Whole-Genome Multilocus Sequence Typing of Extended-Spectrum-Beta-Lactamase-Producing Enterobacteriaceae. J Clin Microbiol. 2016;54(12):2919–2927. doi:10.1128/JCM.01648-16

18. Mikheenko A, Prjibelski A, Saveliev V, Antipov D, Gurevich A. Versatile genome assembly evaluation with QUAST-LG. Bioinformatics. 2018;34(13):i142–i150. doi:10.1093/bioinformatics/bty266

19. Zankari E, Hasman H, Cosentino S, et al. Identification of acquired antimicrobial resistance genes. J Antimicrob Chemother. 2012;67(11):2640–2644. doi:10.1093/jac/dks261

20. Gurevich A, Saveliev V, Vyahhi N, Tesler G. QUAST: Quality assessment tool for genome assemblies. Bioinformatics. 2013;29(8):1072–1075. doi:10.1093/bioinformatics/btt086

21. Wick RR, Schultz MB, Zobel J, Holt KE. Bandage: Interactive visualization of de novo genome assemblies. Bioinformatics. 2015;31(20):3350–3352. doi:10.1093/bioinformatics/btv383

22. García A, Navarro F, Miró E, et al. Acquisition and diffusion of blaCTX-M-9 gene by R478- IncHI2 derivative plasmids. FEMS Microbiol Lett. 2007;271(1):71–77. doi:10.1111/j.1574-6968.2007.00695.x

23. Miró E, Segura C, Navarro F, et al. Spread of plasmids containing the blaVIM-1and blaCTX- Mgenes and the qnr determinant in enterobacter cloacae, klebsiella pneumoniae and klebsiella oxytoca isolates. J Antimicrob Chemother. 2010;65(4):661–665. doi:10.1093/jac/dkp504

24. Nilsen E, Haldorsen BC, Sundsfjord A, et al. Large IncHI2-plasmids encode extended-spectrum β-lactamases (ESBLs) in Enterobacter spp. bloodstream isolates, and support ESBL-transfer to Escherichia coli. Clin Microbiol Infect. 2013;19(11):E516–E518. doi:10.1111/1469-0691.12274

25. Rozwandowicz M, Brouwer MSM, Fischer J, et al. Plasmids carrying antimicrobial resistance genes in Enterobacteriaceae. J Antimicrob Chemother. 2018;73(5):1121–1137. doi:10.1093/jac/dkx488

