## Supplementary figures 1,2. for "Detection of extended-spectrum beta-lactamase (ESBL) genes and plasmid replicons in Enterobacteriaceae using PlasmidSPAdes assembly of short-read sequence data"

**Supplementary figure 1.** Coverage distribution for long contigs (>10kb). Number of contigs for assembly at k-mer size 77. (*median Coverage* = 21,6*,* marked by green line). Yellow lines represent *medianCoverage* times 0,7 and 1,3. Each bar represents all edges with a coverage in a bin of size 1. Blue bars represent long contigs not containing an ESBL gene (**a**) or plasmid replicon (**b**). Orange bars represent long contigs containing an ESBL gene or plasmid replicon.

**a)**

**b)**

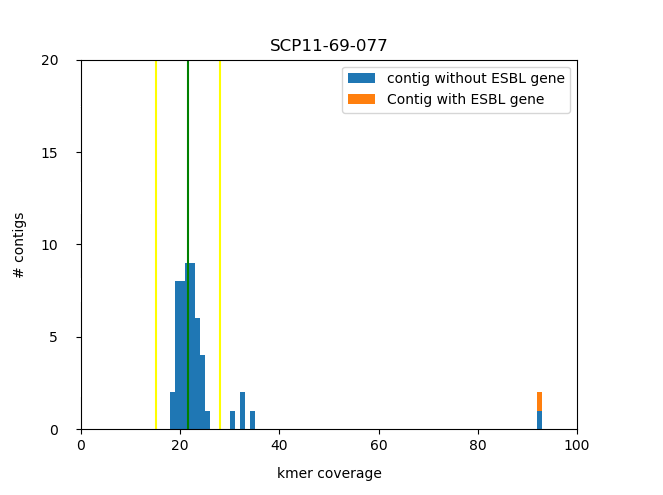

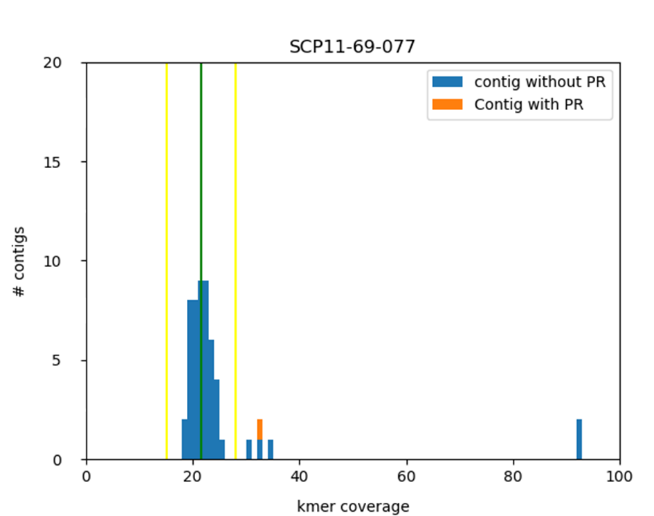

**Supplementary figure 2.** Assembly graph before chromosomal removal were the coverage of the plasmid replicon- and ESBL gene -containing contigs did not deviate from the median coverage of all long edges (>10kb) but were included in the plasmid assembly.

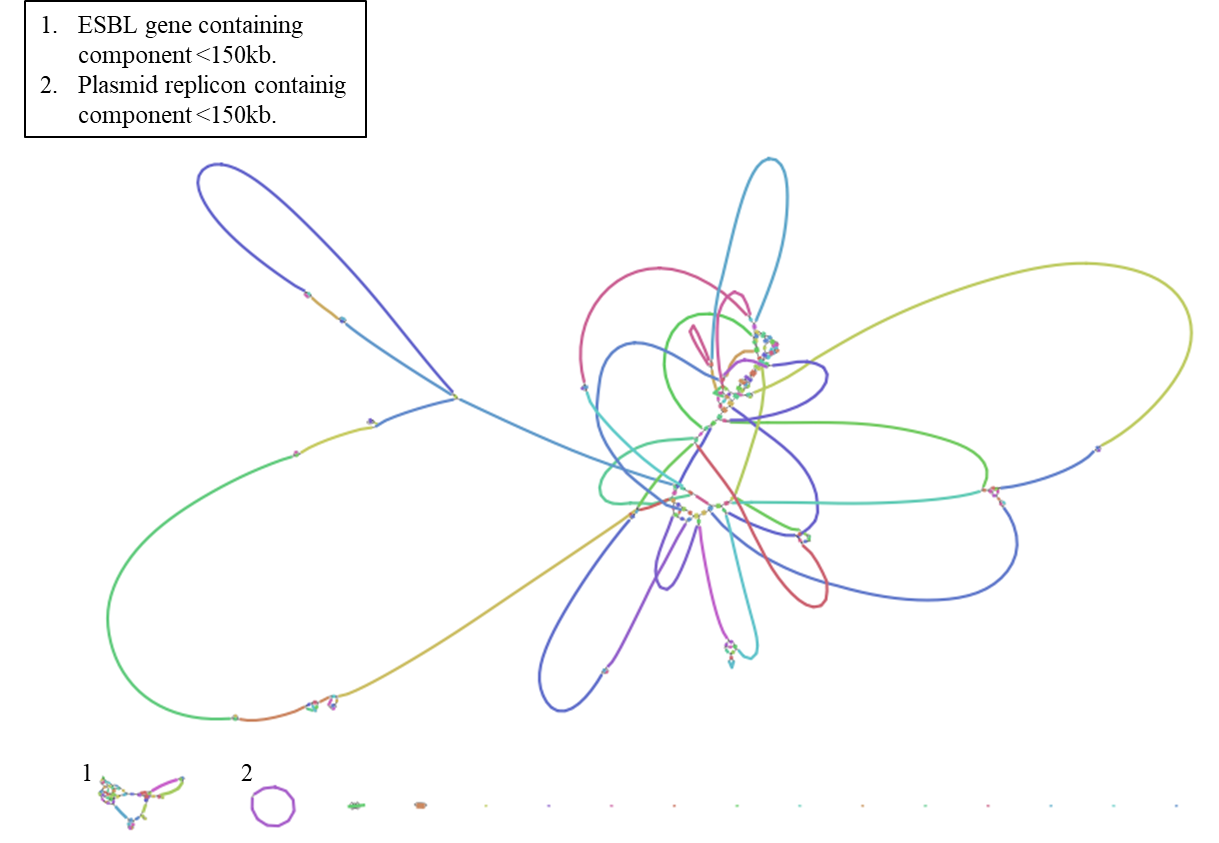
